# Loss of zebrafish *ctnnd2b* results in disorganised forebrain neuron clusters

**DOI:** 10.1101/420828

**Authors:** Wolfgang Hofmeister, Raquel Vaz, Steven Edwards, Alfredo Dueñas Rey, Anna Lindstrand

## Abstract

Delta catenin (CTNND2) is an adhesive junction associated protein belonging to the family of p120ct catenins. It is located on the short arm of chromosome 5, a region deleted in Cri-du-chat syndrome. Heterozygous loss of *CTNND2* function has been linked to autism, schizophrenia, and mild intellectual disability with or without dyslexia-like learning difficulties. To date, most functional studies have focused on homozygous loss of the gene, contradictory to the dominant effect of loss of a single allele observed in neurodevelopmental disorders. Here we show that heterozygous loss of *ctnnd2b* results in a disorganisation and imbalance of neuronal subtypes in forebrain specific regions. Using the zebrafish model, we show that CRISPR/Cas9-induced loss of *ctnnd2b* but not *ctnnd2a* results in an increase in *isl1-*expressing cells and a local reduction of GABA expressing neurons in the optic recess region of the embryonic zebrafish forebrain. Using time-lapse analysis, we found that the disorganised distribution of *is1l-*expressing forebrain neurons was not due to migration defects, but rather an increase in the number of *isl1*-GFP neurons in the optic recess region. Upon closer analysis, these neurons appear disorganised and show an altered morphology and orientation. Overall this data suggests that *ctnnd2* may affect the differentiation cascade of neuronal subtypes in specific regions of the vertebrate brain.

## Introduction

The vertebrate nervous system is a highly organised structure consisting of various neuronal and non-neuronal cell types that need to interact in order to execute complex locomotor, sensory and behavioural tasks. Its function relies on the establishment of a spatial architecture during development which can be divided into gross patterning of the neural tube, proliferation, differentiation and specification of individual neurons, migration to their defined locations in various functional compartments of the brain, dendritic morphogenesis, targeted projection of axons and synapse formation. Neurons are highly diverse and their functional specialisation can be attributed to both their localisation and molecular signature. Broadly, they can be divided into interneurons, relaying signals locally or projection neurons, sending axons to form synapses in other regions of the brain. They are generally further classified into inhibitory and excitatory neurons, characterised by the expression of specific neurotransmitters. Most inhibitory interneurons use the neurotransmitters GABA while most excitatory projecting neurons use Glutamate. Other major neurotransmitters with modulatory actions in the CNS are Acetylcholine, Dopamine, Serotonin, Histamine and Norepinephrine. Understanding the molecular pathways involved in subtype specification, function, and maintenance is of importance.

CTNND2 belongs to the family of p120ct catenins and is an important neural developmental gene which is deleted in Cri-du-chat syndrome. Hemizygous loss of *CTNND2* has been reported to increase the severity of Cri-du-chat (Medina et al., 2000), while isolated heterozygous loss of *CTNND2* function has been linked to autism (Turner et al., 2015), mild intellectual disability with or without dyslexia-like learning difficulties (Hofmeister et al., 2015) (Belcaro et al., 2015), and schizophrenia (Vrijenhoek et al., 2008). During development of the nervous system, the encoded protein ∂-catenin appears to be involved in various neurodevelopment processes. These include neuronal differentiation (Lee et al., 2016), synaptic dysfunction (Israely et al, 2004), dendritic branching, spine morphogenesis (Kim et al., 2008), and maintenance of mature dendrites and dendritic spines (Matter et al., 2009) (Gilbert and Man, 2016). Homozygous mice expressing the GFP-tagged N-terminal domain of CTNND2 show spatial learning difficulties and abnormal fear conditioning (Israely et al., 2004), possibly due to abnormalities in dendritic arbour maintenance (Matter et al., 2009), suggesting that loss of ∂-catenin affects behaviour. However, no early developmental abnormalities were detected and no phenotype was reported in heterozygous mice, contradicting the autosomal dominant role of heterozygous loss of *CTNND2* seen in human patients.

Here we show, using the zebrafish model, that heterozygous loss of *ctnnd2b* but not *ctnnd2a* results in a disorganisation and imbalance of neuronal subtypes in the zebrafish forebrain optic recess region (ORR). Moreover, we observe a local increase in *isl1-*expressing cells and a local reduction of GABA expressing neurons. Using time-lapse analysis, we found that the disorganised distribution of *isl1-*expressing forebrain neurons was not due to migration defects. These clusters of neurons presented an altered morphology and orientation upon closer analysis, suggesting that *ctnnd2* may affect the differentiation cascade of neuronal subtypes in vertebrates.

## Results and Discussion

### *ctnnd2b* LOH but not *ctnnd2a* LOH results in misplaced neurons

The zebrafish genome contains two *ctnnd2* orthologs, due to a genome wide duplication event that occurred during *teleost* evolution. We have previously shown that reduction of *ctnnd2b* expression but not *ctnnd2a* using antisense morpholinos leads to ectopic/misplaced *isl-* expressing neurons in the zebrafish forebrain in the optic recess region (Hofmeister et al., 2014). To further investigate these findings we created stable CRISPR/CAS9 knockout lines in an *isl1:GFP* transgenic background. These lines, named lines 1-4, contain a combination of short deletions and insertions of both *ctnnd2* orthologs, resulting in premature stop codons (refer to methods, supplementary Fig. S1-S4 and Table. S1). To validate the genotype to phenotype penetrance of different allele combinations, double *ctnnd2a* and *ctnnd2b* heterozygotes (line 1 and line 2) were in-crossed and analysed. Heterozygous loss of *ctnnd2b* results in ectopic *isl1*-GFP neurons at 54hpf as previously observed (orange arrows, Fig. 1G and H), compared to wild-type *isl1:GFP* embryos (Fig. 1E), wildtype siblings (Fig. 1F) or embryos with a heterozygous loss of *ctnnd2a* (Fig. S5), confirming our previous finding (Hofmeister et al., 2014). Similar results were seen in mutants with different out of frame deletions in *ctnnd2a* and *ctnnd2b* (Line 2, Fig. S5). The abnormal *isl:GFP* cells are located in the optic recess region (ORR), which can be identified as a distinct morphological entity (Fig. 1A and B) located behind the optic chiasm, rostral to the supraoptic tract (SOT, Fig. 1B) and between the anterior commissure (AC) of the telencephalon and the post-optic commissure (POC) of the diencephalon which gives rise to the preoptic area (Affaticati et al., 2015). Although some *isl1-*positive neurons are occasionally seen in the optic recess region of wildtype *isl1:GFP* animals (Fig. 1I, Fig. S7, Fig. 3D), the GFP-positive population is significantly expanded in knockout fish both in number and intensity (Fig. 1E-M, Fig. 3D’). Confocal analysis at a higher magnification shows that the ectopic neurons still project axons into the post-optic commissure (Fig. 1N-P’). However, the neurons display a vertical rather than horizontal orientation and the cluster seems disorganised compared to neurons in wildtype embryos that form part of the diencephalic population (Fig. 1N-P’). Ectopic neurons are also still present at later stages (6dpf) analysed (Fig. 1L, M).

**Fig 1.**
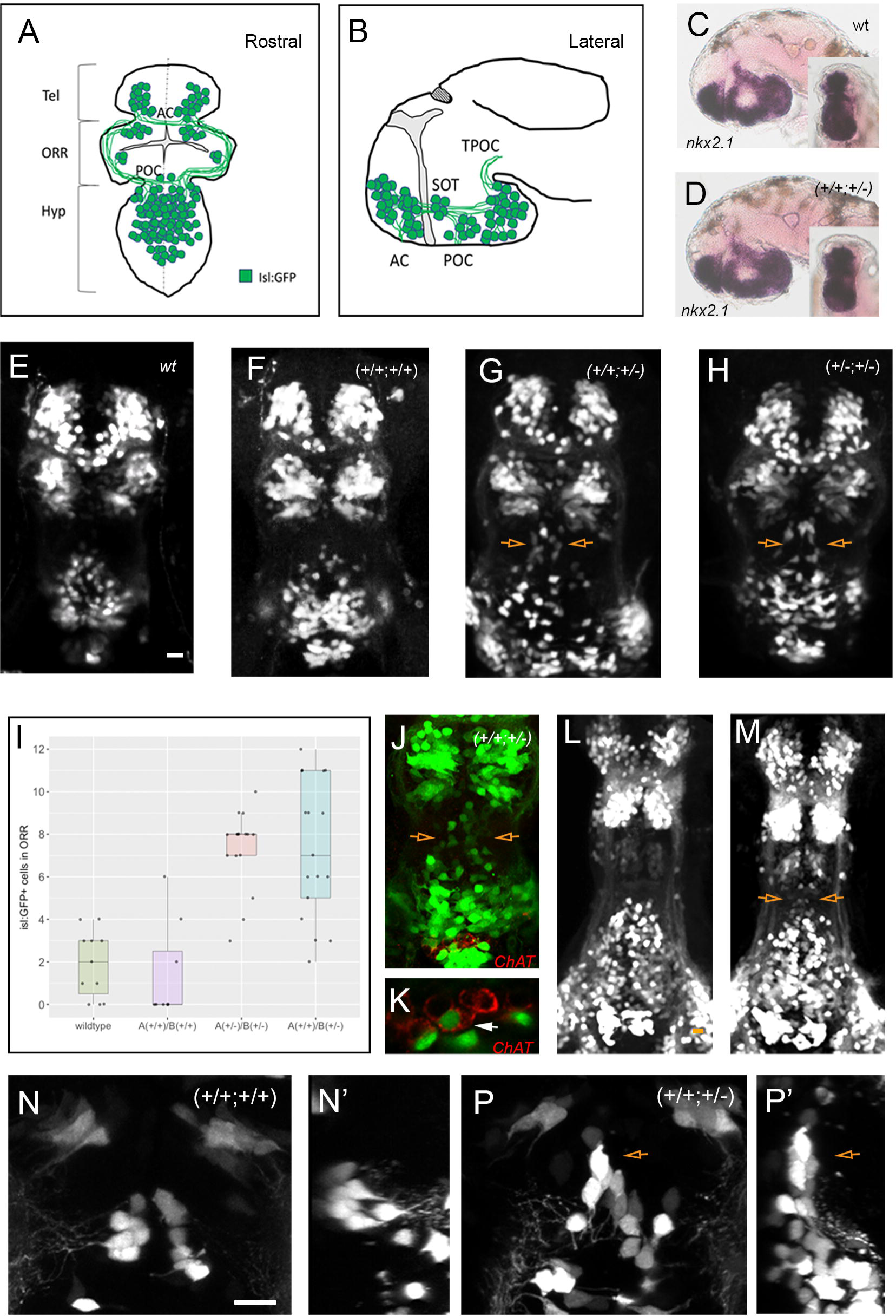
Heterozygous loss of *ctnnd2b* results in ectopic neuron specification and abnormal positioning. Schematic showing *isl:GFP* neuron population in the zebrafish forebrain at 2dpf (A, B). Anterior view in (A) and lateral view in (B) with rostral to the left. Confocal z-stacks of zebrafish embryos taken from an anterior view are shown (E-H and J-M). 54hpf embryo brains are shown in (E-H) and 6dpf brain in (L and M). Brightfield *in situ* images of laterally mounted 54hpf brains stained with a riboprobe for *nkx2.1* are shown in (C, D) with anterior views in the inserts. No difference in nkx2.1 staining was observed. Wildtype *isl:GFP* transgenic zebrafish are shown in (C and E). Non-carrier sibling (wildtype larvae) from a cross of a double *ctnnd2a* and *ctnnd2b* heterozygotes are shown in (F, L, N and N’), a heterozygous *ctnnd2b* carrier in (G, J, K, M, P and P’) and a double *ctnnd2a* and *ctnnd2b* heterozygote in (H). Count of misplaced *isl:GFP* cells at the optic recess region is shown in (I) with heterozygotes (A(+/+)/B(+/-); n=17) and double heterozygotes (A(+/-)/B(+/-); n=16) significantly increased in number (Wilcoxon test; p<0.001) when compared to non-carrier siblings (A(+/+)/B(+/+);n=8) or non-sibling *isl1:GFP* embryos (n=11) (I). Confocal optical slices at the level of ectopic neurons is shown in N and P from an anterior view with lateral projections in N’ and P’ with rostral to the right. Scale bar =10μm

### Ectopic *isl:GFP* neurons do not express ChAT in early development

In mammals, ISL1 has been shown to form a complex with LHX3 to promote cholinergic gene expression in basal forebrain cholinergic neurons (Hyong Ho Cho et al., 2013). This neuronal population is involved in attention, memory, reward pathways, and motor activity (for review see (Allaway and Machold, 2017). Forebrain cholinergic neurons have an embryonic origin in the ventral telencephalon (subpallium), in a region that expresses the transcription factor *nkx2.1*. In mammals the *nkx2.1* expression domain encompasses the medial ganglionic eminences (MGE), septum, and post optic area. The equivalent *nkx2.1* expression domain in zebrafish where the *isl1* forebrain cells reside encompasses the ventral telencephalon, optic recess region, ventral diencephalon, and hypothalamus (Manoli and Driever, 2014). *In situ* analysis revealed no difference in patterning of this *nkx2.1* region between wildtype and heterozygous embryos (Fig. 1C, D) suggesting normal patterning of this region. In order to assess whether zebrafish *isl1* forebrain neurons also express acetyl choline we used an antibody against choline acetyl transferase (ChAT). However no staining was evident in the majority forebrain *isl:GFP* neurons at 54hpf (Fig. 1J). Instead CHAT staining at 54hpf is confined to a cluster of neurons in the hypothalamus (Fig. 1J and K), some of which but not all express *isl1* at the specific stage imaged (Fig. 1J). In *drosophila* loss of *isl1* results in loss of dopamine and serotonin synthesis (Thor and Thomas, 1997). Despite ectopic *isl:GFP* expression in neurons we see no change of dopaminergic, or serotoninergic cells in the zebrafish forebrain as assessed by TH1 and 5-HT staining respectively, and there is no overlap between *isl1:GFP* expression and TH1 or 5-HT in any of the misplaced cells at 2dpf (supplementary Fig. S6A-B and E-G).

### *Ctnnd2b LOH* results in abnormal GABAnergic neurogenesis

The ventral telencephalon is also the birthplace of GABAergic neurons. The choice as to whether *nkx2.1* expressing cells in the ventral telencephalon become GABAergic or cholinergic is due to a distinct expression of a combination of transcription factors. Post mitotic neurons fated to become GABAergic neurons express high levels of *lhx6,* a transcription factor essential for differentiation to GABAergic neurons (Liodis et al., 2007). Conversely, in post-mitotic *nkx2.1* positive neurons expressing high levels of *isl1* and *lhx8,* the expression of *lhx6* is supressed and these neurons become forebrain cholinergic neurons (Cho et al., 2014). Thus, the differentiation into GABAergic neurons is inhibited by the expression of *isl1* (Hutchison 2006). We therefore decided to assess the expression GABAergic neurotransmitters. Staining with an antibody against GABA shows that GABAergic neurons in diencephalon and ORR are intermingled but rarely overlapping with *isl1:GFP* as expected (Fig. 2A,B,D,E). Conversely, ectopic *isl1* neurons in regions were GABA is normally expressed do not show GABA expression, and populations of GABA stained neurons appear to be more loosely packed together in both heterozygous *ctnnd2b* or double heterozygous embryos then in controls (compare Fig. 2A and D to B and E). A similar result is seen in double homozygous embryos (supplementary Fig. S6H). Resliced side projections confirm the presence of ectopic *isl*-GFP cells in the normally GABA+ ORR (compare Fig. S6I and J). Count of GABAergic neurons throughout the Z-sections taken at the ORR region confirmed a reduction in GABA+ neurons in heterozygous *ctnnd2b* embryos in addition to the disorganisation (Fig. 2C).

**Fig 2.**
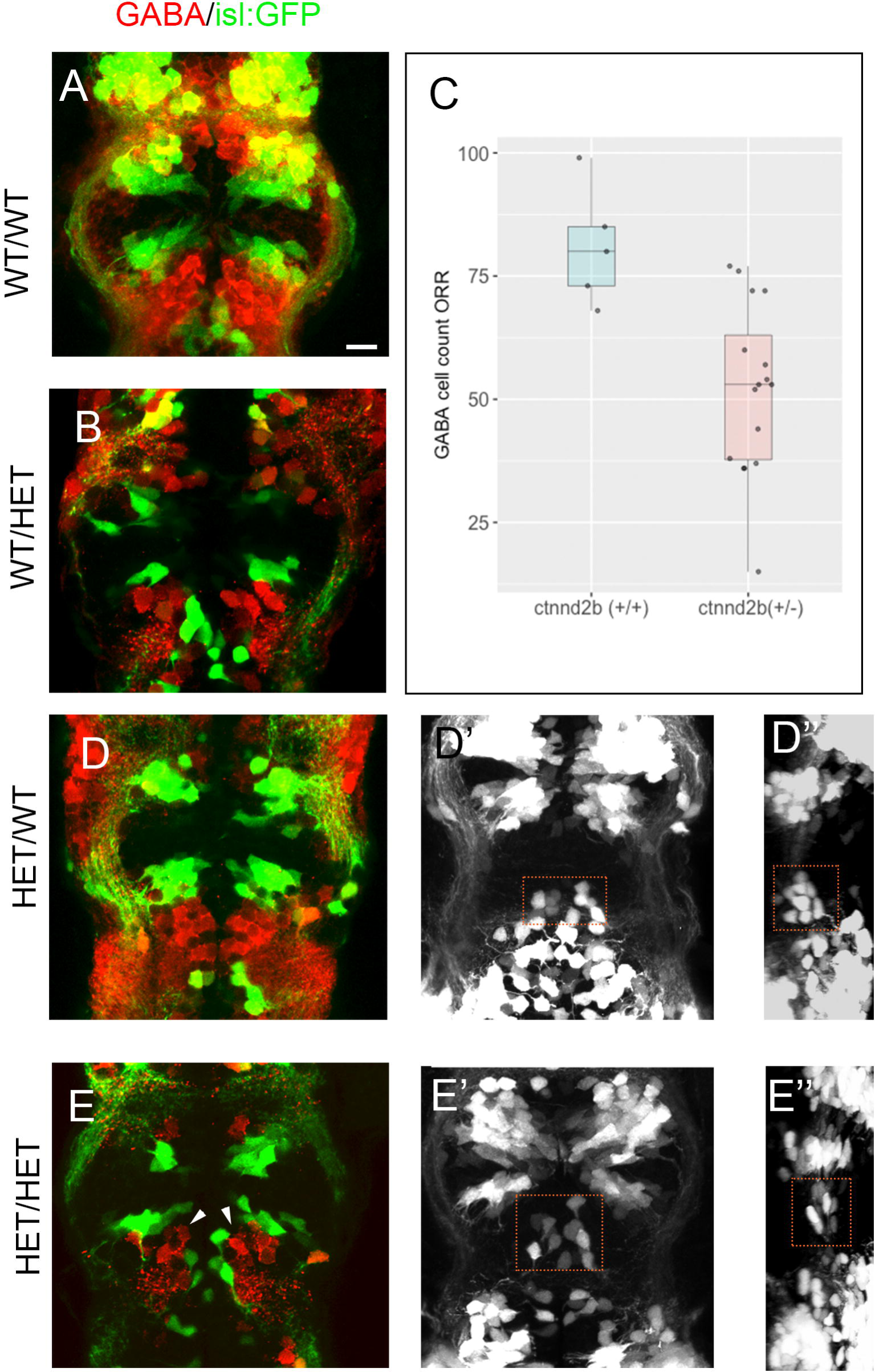
CTNND2B LOH results in disorganised GABA neurons. Confocal optical slices at the level of ectopic *isl:GFP* neurons (green), showing GABA stained cells (red) in (A,B,D,E). Equivalent confocal z-stack of (D) and (E) showing the positioning of normal *isl:GFP* cluster (D’) and abnormal ectopic cells (E’) is shown and the lateral view of the positioning of equivalent ectopic *isl:GFP* neuron cluster in (E’’) and (B’’’). anterior views are shown in (A, B, D, D’, E, E’) and resliced Lateral views in (D’’ and E’’). Count of anti-GABA stained cell throughout the ORR is shown in (C). Where the optic recess region was defined as the region from the POC where we start seeing ectopic *isl1:GFP* cells to the start of *isl:GFP* cluster of cells branching from the SOT. A significant reduction is observed (Wilcoxon test, p<0.01) when comparing siblings with wildtype *ctnnd2b* (n=5) compared to those heterozygous loss of *ctnnd2b* (n=16) irrespective of *ctnnd2a* genotype. Scale bar =10μm

### Increased *isl1* expressing cells at optic recess region due to specification not migration

In order to asses if the observed phenotype was due to migration or abnormal specification we imaged embryos between 32 and 48 hpf. Staging was additionally assessed by growth of visible axon tracts such as the trigeminal and post-optic commissure. Initially the number of *GFP* expressing cells were similar between *isl:GFP* in-crossed control larvae (Fig. 3B and Movie S7) and in animals missing one functional copy of *ctnnd2b* (Fig. 3B’ and Movie S7). However, as development proceeds we see a decrease in cells expressing *GFP* in controls, with only a few cells remaining at the end of the recording period 48hpf (Fig. 3D and Movie. S7). In contrast, in mutant embryos we observe an increase in the number of GFP-positive cells arising in the optic recess region, with no indication of downregulation of *GFP* throughout the developmental stages analysed (Fig. 3D’ and Movie. S7). In addition, *GFP*-positive cells appear less organised, supporting the observations from fixed embryos imaged using confocal microscopy at a slightly later stage at 54hpf (Fig. 1).

**Fig 3.**
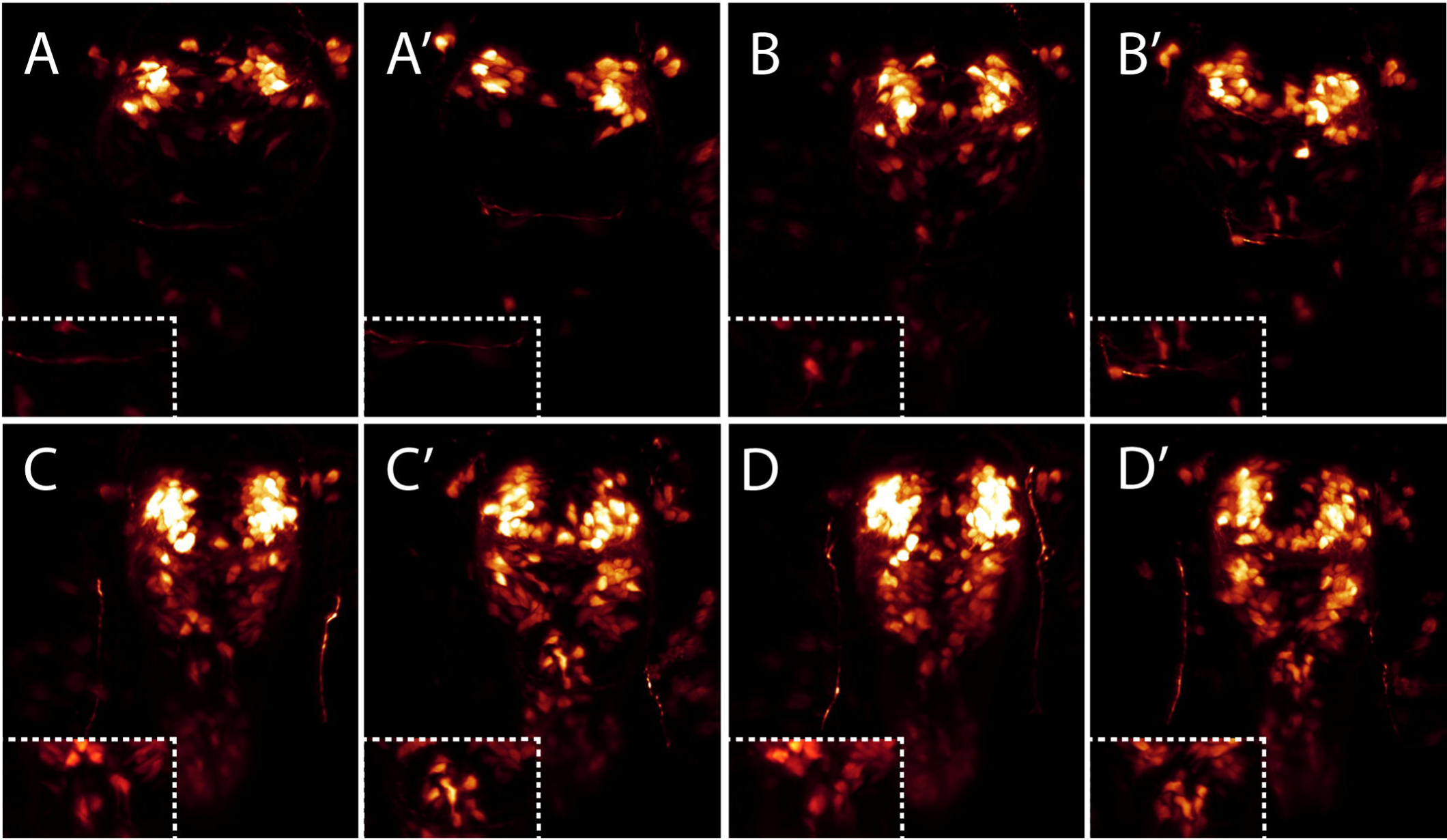
Increase in *isl1* positive cells is due to miss-specification not migration. 3D renders of timelapse imaged wildtype (A-D) and abnormal *ctnnd2b* heterozygous embryos (A’-D’) with highlighted ORR (insets). Embryos at 32hpf prior to appearance of neurons at ORR are shown in (A). Similarly, numbers of neurons start appearing at the optic recess region by 37hpf (B) and 42.5hpf (C). By the end of recording at approx. 48hpf only a few *isl1:GFP* cells are evident in wildtype embryos compared to heterozygous embryos (D).

In summary our results show that knockout of *ctnnd2* results in in the overexpression of *isl1* positive cells at the optic recess region which are highly disorganised and either directly or indirectly affect the local differentiation and organisation of GABA neurons in that region. Recently *CTNND2* has shown to be a target gene of REST and CoREST (Abrajano et al., 2009), transcription factors which are thought to modulate neuronal subtype specification (Lee et al., 2016). Our study supports a role for ∂-catenin in subtype specification. Studies have implicated ∂-catenin as a modulator of canonical Wnt and RhoA signalling as well as modulating Cadherin stability. However, its exact role in these pathways is still a matter of debate. The transcription factor ISL1 is known to be downstream of canonical WNT signalling in cardiac development (Qyang et al., 2007)(Lu et al., 2014). Whether loss of CTNND2 causes abnormal ISL1 /GABA neuronal balance in specific regions of the mammalian brain and if it acts through an increase in canonical WNT signalling is yet to be determined.

*CTNND2* haploinsufficiency has been linked to autism and mild neurodevelopmental disorders specifically reading difficulties. Reduced GABAergic action has been observed in the brain of autistic individuals (Fatemi et al., 2009)(Robertson et al., 2016) (Oblak et al., 2011) and genes encoding GABA receptor subunits have been associated with autism in linkage and copy number studies (Ma et al., 2005) suggesting imbalance or dysfunction of the GABAergic system a potential cause of autism. Our study supports a role for CTNND2 haploinsufficiency in the aetiology of previously associated neural developmental disorders due to abnormal GABA neurons differentiation in specific brain regions.

## Materials and Methods

### Zebrafish husbandry

All fish were maintained on a 14 h day/night cycle at the Karolinska Institute zebrafish core facility, located in the department of Comparative Medicine. Embryos were produced by light-induced spawning. Tg(isl1:GFP)rw0 transgenic strains (*isl:GFP* for short) or TL strains were used for all experiments.

### CRISPR/Cas9-mediated knockout of *ctnnd2* orthologues

Zebrafish mutant lines were established using a protocol adapted from Varshney *et al* (2015). sgRNAs and Cas9 protein mix was injected into 1-cell embryos and efficiency was tested using the PCR-STAT method. The target sequence of active sgRNAs were in exon 5 of *ctnnd2a* (5’-GGT TCA GCG AGC GAT ACA CC-3’) and exon 9 of *ctnnd2b* (5’-GGT GGA AGC GCC CGG GCC AG-3’). Primers used for screening were *ctnnd2a-F (5’-*TCA CTC TCC TTA CAG GTG ACA-3’) and *ctnnd2a-R* (5’-GGT TGC GTA GCC AGG TGT AT-3’) or *ctnnd2b-F* (5’-CCA GGT CCT GAA CTT TCT GC-3’) and *ctnnd2b –R* (5’-GGT CCT TCT GGA TGC TGT CC-3’), respectively. Embryos were raised and founders identified using the same PCR-STAT method. Founders were outcrossed to wildtype or i*sl:GFP* fish and raised to adult stage. Heterozygous F1 adults were identified using fin clipping and subsequent Sanger sequencing.

### Characterisation of mutant lines

We identified a total of 2 lesions for *ctnnd2a*, an 1bp deletion and an 8bp deletion. For *ctnnd2b* we identified a total of three, a 7bp deletion, a combined *6bp* deletion and 10bp insertion, and a 1bp deletion, the latter was not used for further analysis. All of these lesions were predicted to cause a frameshift resulting in premature termination of the protein sequence (Fig. S1-4) Combinations of these resulted in five distinct F1 carrier lines., Line1; 8bp deletion in ctnnd2a and 6bp deletion/10bp insertion in ctnnd2b, Line2; 1bp deletion in ctnnd2a and 7bp deletion in ctnnd2b, Line3; with only the 7bp deletion in *ctnnd2b,* and Line 4 a with only the 6bp deletion/10bp insertion in ctnnd2b (Table. S1). F1 fish were incrossed or outcrossed and larvae analysed for phenotypes as indicated throughout the manuscript. Embryos were fixed and phenotyped via confocal microscopy before being genotyped by sanger sequencing. Alternatively, larvae were anesthetized and their tails were cut to allow genotyping before tissue fixation and image analysis

### Immunofluorescence and *in situ* hybridisations

Embryos were fixed in 4% (w/v) paraformaldehyde (PFA) overnight at 4°C. Whole larvae or dissected brains were processed for immunofluorescence using standard protocol (Devine et al., 2003). Primary antibodies used were rabbit anti-Th1 (cat# P40101; Peel-freez biologicals), mouse anti-acetylated tubulin (cat# T6793; Sigma), rabbit anti-GABA (cat# A2052; Sigma), goat anti-CHAT (cat# AB144P Sigma/Merck), rabbit anti-Serotonin (cat#S5545; Sigma). Secondary antibodies used were goat anti-mouse Alexa 594 (cat # A11032 Life Technologies, Carlsbad, California, USA), donkey anti-goat Alexa 546 (cat# A11056 Life technologies). *In situ* staining was performed as previously described (Hjorth *et al.,* 2001). The *nkx2.1* riboprobe was synthesised using the DIG labelling kit (Roche) using the following PCR template with a T7 RNA polymerase site attached to the 5’ end of the reverse primer (forward primer: 5’-TTG CCT CGT ACA GAC AAC CC-3’, reverse primer: 5’-<u>TAA TAC GAC TCA CTA TAG GGG</u> TCA CCA CGT CCT GCC ATA A-3’). Embryos were mounted in gelvatol for imaging.

### Confocal imaging and Data analysis

Transgenic embryos fixed in 4% PFA or immunostained embryos were mounted rostrally or laterally in gelvatol mounting medium and imaged using a Zeiss LSM 700,800 or Olympus FV1000. Images where blinded and *isl:GFP* or GABA stained cells counted throughout individual confocal sections. Cell counts were subsequently corelated back to the genotype of the embryos as identified by sanger sequencing. ISL1-positive cell count from multiple experiments/crosses were pooled together (supplementary table S2). For GABA-positive cell counts, data was obtained from two separate crosses of double heterozygous adults and one cross of double homozygous adults (table S3). *dplyr* package in R-studio was used for data handling, statistical analysis and visualisation using the *ggplot2* package. The Mann-whitney-wilcoxon tests was used to compare between samples.

### Lightsheet microscopy

31hpf embryos were anesthetised in tricaine (MS222) and mounted in 1% low melting point agarose containing 1:2000 F-Z 050 fluorescent microspheres (Merck) in a glass capillary. The agarose cylinder was extruded into a sample chamber containing E3 medium at 28.5°C. Two channel images were acquired using a Light Sheet Z.1(Carl Zeiss, Germany) with a water dipping 20X detection objective (W-Plan-APOCHROMAT-1.0NA) and dual side 10X illumination objectives (LSFM, 0.2NA). Samples were illuminated from two sides and Z-stacks were acquired from 3 angles (covering 90°) every 20 min for 16 hours. Data from multiple angles was registered and deconvolved in the Multiview Reconstruction Fiji plugin (Preibisch et al., 2010). Briefly, beads were detected in one channel and used to register views and correct drift. Transformations were duplicated to the GFP channel and images were fused by multi-view deconvolution, performed on the GPU (Preibisch et al., 2014). Deconvolved images were rendered and movies created in Imaris 9.0 (Bitplane).

